# The genetic structure of the world’s first farmers

**DOI:** 10.1101/059311

**Authors:** Iosif Lazaridis, Dani Nadel, Gary Rollefson, Deborah C. Merrett, Nadin Rohland, Swapan Mallick, Daniel Fernandes, Mario Novak, Beatriz Gamarra, Kendra Sirak, Sarah Connell, Kristin Stewardson, Eadaoin Harney, Qiaomei Fu, Gloria Gonzalez-Fortes, Songül Alpaslan Roodenberg, György Lengyel, Fanny Bocquentin, Boris Gasparian, Janet M. Monge, Michael Gregg, Vered Eshed, Ahuva-Sivan Mizrahi, Christopher Meiklejohn, Fokke Gerritsen, Luminita Bejenaru, Matthias Blueher, Archie Campbell, Gianpero Cavalleri, David Comas, Philippe Froguel, Edmund Gilbert, Shona M. Kerr, Peter Kovacs, Johannes Krause, Darren McGettigan, Michael Merrigan, D. Andrew Merriwether, Seamus O’Reilly, Martin B. Richards, Ornella Semino, Michel Shamoon-Pour, Gheorghe Stefanescu, Michael Stumvoll, Anke Tönjes, Antonio Torroni, James F. Wilson, Loic Yengo, Nelli A. Hovhannisyan, Nick Patterson, Ron Pinhasi, David Reich

## Abstract

We report genome-wide ancient DNA from 44 ancient Near Easterners ranging in time between ~12,000-1,400 BCE, from Natufian hunter-gatherers to Bronze Age farmers. We show that the earliest populations of the Near East derived around half their ancestry from a ‘Basal Eurasian’ lineage that had little if any Neanderthal admixture and that separated from other non-African lineages prior to their separation from each other. The first farmers of the southern Levant (Israel and Jordan) and Zagros Mountains (Iran) were strongly genetically differentiated, and each descended from local hunter-gatherers. By the time of the Bronze Age, these two populations and Anatolian-related farmers had mixed with each other and with the hunter-gatherers of Europe to drastically reduce genetic differentiation. The impact of the Near Eastern farmers extended beyond the Near East: farmers related to those of Anatolia spread westward into Europe; farmers related to those of the Levant spread southward into East Africa; farmers related to those from Iran spread northward into the Eurasian steppe; and people related to both the early farmers of Iran and to the pastoralists of the Eurasian steppe spread eastward into South Asia.

---

Between 10,000-9,000 BCE, humans began practicing agriculture in the Near East^1^. In the ensuing five millennia, plants and animals domesticated in the Near East spread throughout West Eurasia (a vast region that also includes Europe) and beyond. The relative homogeneity of present-day West Eurasians in a world context^2^ suggests the possibility of extensive migration and admixture that homogenized geographically and genetically disparate sources of ancestry. The spread of the world’s first farmers from the Near East would have been a mechanism for such homogenization. To date, however, due to the poor preservation of DNA in warm climates, it has been impossible to study the population structure and history of the first farmers and to trace their contribution to later populations.

In order to overcome the obstacle of poor DNA preservation, we took advantage of two methodological developments. First, we sampled from the inner ear region of the petrous bone^3,4^ that can yield up to ~100 times more endogenous DNA than other skeletal elements^4^. Second, we used in-solution hybridization^5^ to enrich extracted DNA for about 1.2 million single nucleotide polymorphism (SNP) targets^6,7^, making efficient sequencing practical by filtering out microbial and non-informative human DNA. We merged all sequences extracted from each individual, and randomly sampled a single sequence to represent each SNP, restricting to individuals with at least 9,000 SNPs covered at least once. We obtained genome-wide data passing quality control for 45 individuals on whom we had a median coverage of 172,819 SNPs (Methods). We assembled radiocarbon dates for 26 individuals (22 new generated for this study) (Supplementary Data Table 1).

The newly reported ancient individuals date to ~12,000-1,400 BCE and come from the southern Caucasus (Armenia), northwestern Anatolia (Turkey), Iran, and the southern Levant (Israel and Jordan) (Supplementary Data Table 1, Fig. 1a). (One individual had a radiocarbon date that was not in agreement with the date of its archaeological context and was also a genetic outlier.) The samples include Epipaleolithic Natufian hunter-gatherers from Raqefet Cave in the Levant (12,000-9,800 BCE); a likely Mesolithic individual from Hotu Cave in the Alborz mountains of Iran (probable date of 9,100-8,600 BCE); Pre-Pottery Neolithic farmers from ‘Ain Ghazal and Motza in the southern Levant (8,300-6,700 BCE); and early farmers from Ganj Dareh in the Zagros mountains of western Iran (8,200-7,600 BCE). The samples also include later Neolithic, Chalcolithic (~4,800-3,700 BCE), and Bronze Age (~3,350-1,400 BCE) individuals (Supplementary Information, section 1). We combined our data with previously published ancient data^7,8,9,10,8,10–15^ to form a dataset of 281 ancient individuals. We then further merged with 2,583 present-day people genotyped on the Affymetrix Human Origins array^13,16^ (238 new) (Supplementary Data Table 2; Supplementary Information, section 2). We grouped the ancient individuals based on archaeological culture and chronology (Fig. 1a; Supplementary Data Table 1). We refined the grouping based on patterns evident in Principal Components Analysis (PCA)^17^ (Fig. 1b; Extended Data Fig. 1), ADMIXTURE model-based clustering^18^ (Fig. 1c), and ‘outgroup’ *f*_3_-analysis (Extended Data Fig. 2). We used *f*_4_-statistics to identify outlier individuals and to cluster phylogenetically indistinguishable groups into ‘Analysis Labels’ (Supplementary Information, section 3).

**Figure 1:**
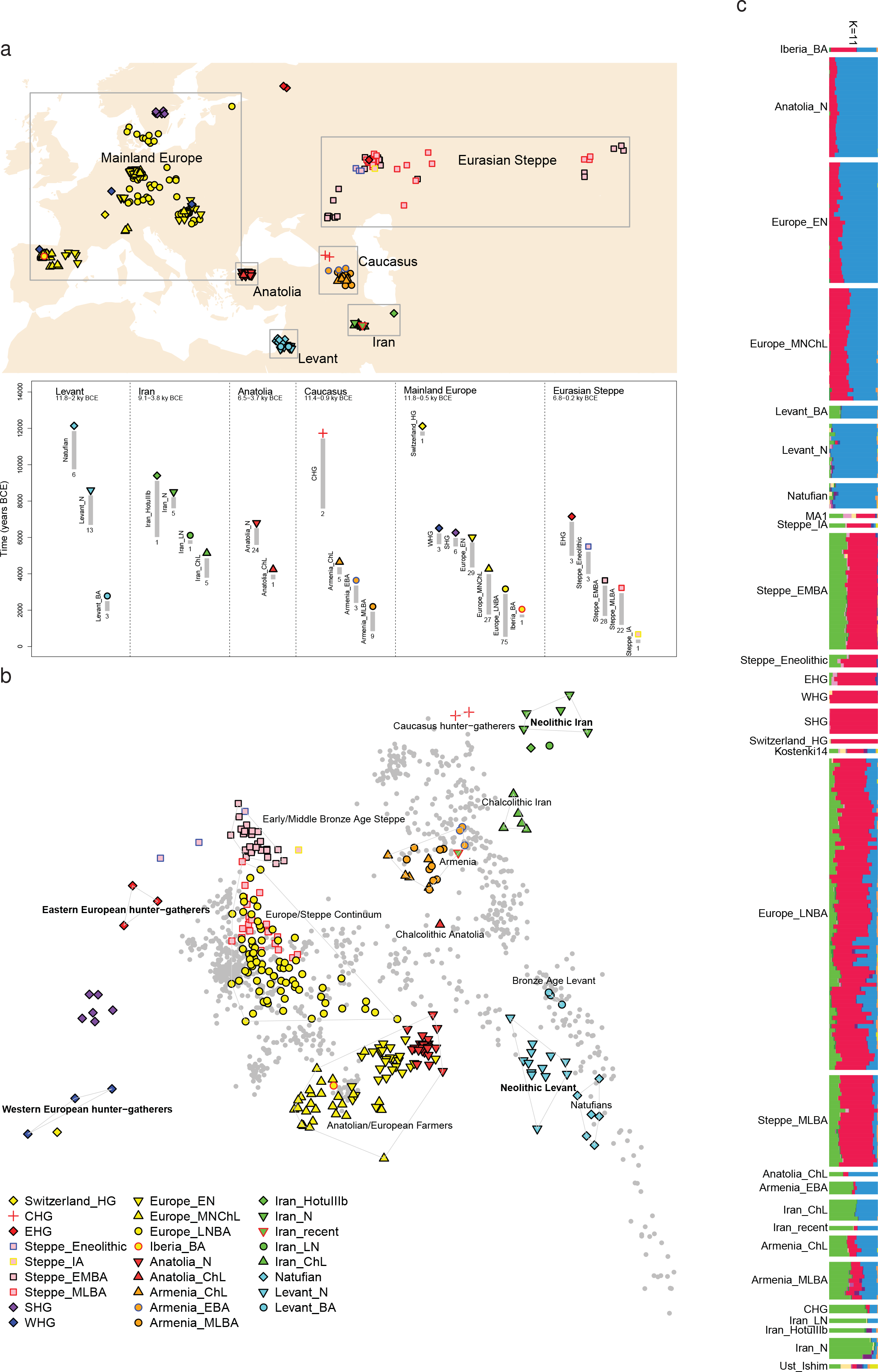
Genetic structure of ancient West Eurasia. (a) Sampling locations and times in six West Eurasian regions. Sample sizes for each population are given below each bar. Abbreviations used: E: Early, M: Middle, L: Late, HG: Hunter-Gatherer, N: Neolithic, ChL: Chalcolithic, BA: Bronze Age, IA: Iron Age. (b) Principal components analysis of 991 present-day West Eurasians (grey points) with 278 projected ancient samples (excluding the Upper Paleolithic Ust_Ishim, Kostenki14, and MA1). To avoid visual clutter, population labels of present-day individuals are shown in Extended Data Fig. 1. (c) ADMIXTURE model-based clustering analysis of 2,583 present-day humans and 281 ancient samples; we show the results only for ancient samples for K=11 clusters.

We analyzed these data to address six questions. (1) Previous work has shown that the first European farmers harboured ancestry from a Basal Eurasian lineage that diverged from the ancestors of north Eurasian hunter-gatherers and East Asians before they separated from each other^13^ What was the distribution of Basal Eurasian ancestry in the ancient Near East? (2) Were the first farmers of the Near East part of a single homogeneous population, or were they regionally differentiated? (3) Was there continuity between late pre-agricultural hunter-gatherers and early farming populations, or were the hunter-gatherers largely displaced by a single expansive population as in early Neolithic Europe?^8^ (4) What is the genetic contribution of these early Near Eastern farmers to later populations of the Near East? (5) What is the genetic contribution of the early Near Eastern farmers to later populations of mainland Europe, the Eurasian steppe, and to populations outside West Eurasia? (6) Do our data provide broader insights about population transformations in West Eurasia?

## Basal Eurasian ancestry was pervasive in the ancient Near East and associated with reduced Neanderthal ancestry

The ‘Basal Eurasians’ are a lineage hypothesized^13^ to have split off prior to the differentiation of all other Eurasian lineages, including both eastern non-African populations like the Han Chinese, and even the early diverged lineage represented by the genome sequence of the ~45,000 year old Upper Paleolithic Siberian from Ust’-Ishim^11^. To test for Basal Eurasian ancestry, we computed the statistic *f_4_(Test*, Han; Ust’-Ishim, Chimp) (Supplementary Information, section 4), which measures the excess of allele sharing of Ust’-Ishim with a variety of *Test* populations compared to Han as a baseline. This statistic is significantly negative (Z<-3.7) for all ancient Near Easterners as well as Neolithic and later Europeans, consistent with their having ancestry from a deeply divergent Eurasian lineage that separated from the ancestors of most Eurasians prior to the separation of Han and Ust’-Ishim. We used *qpAdm*^7^ to estimate Basal Eurasian ancestry in each *Test* population. We obtain the highest estimates in the earliest populations from both Iran (66±13% in the likely Mesolithic sample, 48±6% in Neolithic samples), and the Levant (44±8% in Epipaleolithic Natufians) (Fig. 2), showing that Basal Eurasian ancestry was widespread across the ancient Near East.

**Figure 2:**
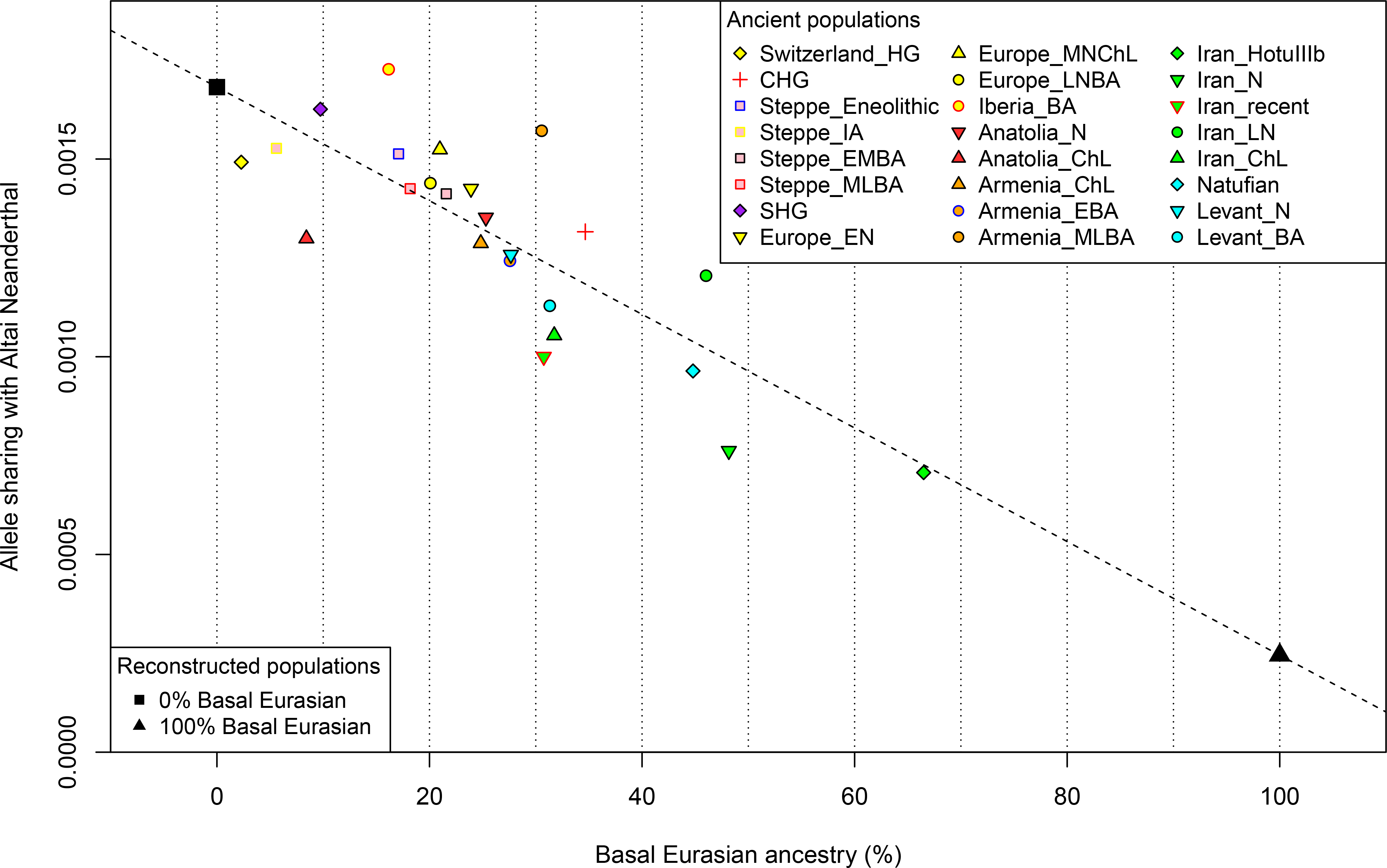
Basal Eurasian ancestry explains the reduced Neanderthal admixture in West Eurasians. Basal Eurasian ancestry estimates are negatively correlated to a statistic measuring Neanderthal ancestry *f*_4_(*Test*, Mbuti; Altai, Denisovan).

West Eurasians harbour significantly less Neanderthal ancestry than East Asians ^19,20–23^, which could be explained if West Eurasians (but not East Asians) have partial ancestry from a source diluting their Neandertal inheritance^21^. Supporting this theory, we observe a negative correlation between Basal Eurasian ancestry and the rate of shared alleles with Neanderthals^19^ (Supplementary Information, section 5; Fig. 2). By extrapolation, we infer that the Basal Eurasian population had lower Neanderthal ancestry than non-Basal Eurasian populations and possibly none (ninety-five percent confidence interval truncated at zero of 0-60%; Fig. 2; Methods). The finding of little if any Neanderthal ancestry in Basal Eurasians could be explained if the Neanderthal admixture into modern humans 50,000-60,000 years ago^11^ largely occurred after the splitting of the Basal Eurasians from other non-Africans.

It is striking that the highest estimates of Basal Eurasian ancestry are from the Near East, given the hypothesis that it was there that most admixture between Neanderthals and modern humans occurred^19, 24^. This could be explained if Basal Eurasians thoroughly admixed into the Near East before the time of the samples we analyzed but after the Neanderthal admixture. Alternatively, the ancestors of Basal Eurasians may have always lived in the Near East, but the lineage of which they were a part did not participate in the Neanderthal admixture.

A population without Neanderthal admixture, basal to other Eurasians, may have plausibly lived in Africa. Craniometric analyses have suggested that the Natufians may have migrated from north or sub-Saharan Africa^25,26^, a result that finds some support from Y chromosome analysis which shows that the Natufians and successor Levantine Neolithic populations carried haplogroup E, of likely ultimate African origin, which has not been detected in other ancient males from West Eurasia (Supplementary Information, section 6) ^7,8^. However, no affinity of Natufians to sub-Saharan Africans is evident in our genome-wide analysis, as present-day sub-Saharan Africans do not share more alleles with Natufians than with other ancient Eurasians (Extended Data Table 1). (We could not test for a link to present-day North Africans, who owe most of their ancestry to back-migration from Eurasia^27,28^.) The idea of Natufians as a vector for the movement of Basal Eurasian ancestry into the Near East is also not supported by our data, as the Basal Eurasian ancestry in the Natufians (44±8%) is consistent with stemming from the same population as that in the Neolithic and Mesolithic populations of Iran, and is not greater than in those populations (Supplementary Information, section 4). Further insight into the origins and legacy of the Natufians could come from comparison to Natufians from additional sites, and to ancient DNA from north Africa.

## Extreme regional differentiation in the ancient Near East

PCA on present-day West Eurasian populations (Methods) (Extended Data Fig. 1) on which we projected the ancient individuals (Fig. 1b) replicates previous findings of a Europe-Near East contrast along the horizontal Principal Component 1 (PC1) and parallel clines (PC2) in both Europe and the Near East (Extended Data Fig. 1)^7,8,13^. Ancient samples from the Levant project at one end of the Near Eastern cline, and ancient samples from Iran at the other. The two Caucasus Hunter Gatherers (CHG)^9^ are less extreme along PC1 than the Mesolithic and Neolithic individuals from Iran, while individuals from Chalcolithic Anatolia, Iran, and Armenia, and Bronze Age Armenia occupy intermediate positions. Qualitatively, the PCA has the appearance of a quadrangle whose four corners are some of the oldest samples: bottom-left: Western Hunter Gatherers (WHG), top-left: Eastern Hunter Gatherers (EHG), bottom-right: Neolithic Levant and Natufians, top-right: Neolithic Iran. This suggests the hypothesis that diverse ancient West Eurasians can be modelled as mixtures of as few as four streams of ancestry related to these populations, which we confirmed using *qpWave*^7^ (Supplementary Information, section 7).

We computed squared allele frequency differentiation between all pairs of ancient West Eurasians^29^ (Methods; Fig. 3; Extended Data Fig. 3), and found that the populations at the four corners of the quadrangle had differentiation of F_ST_=0.08−0.15, comparable to the value of 0.09−0.13 seen between present-day West Eurasians and East Asians (Han) (Supplementary Data Table 3). In contrast, by the Bronze Age, genetic differentiation between pairs of West Eurasian populations had reached its present-day low levels (Fig. 3): today, F_ST_ is ≤0.025 for 95% of the pairs of West Eurasian populations and ≤0.046 for all pairs. These results point to a demographic process that established high differentiation across West Eurasia and then reduced this differentiation over time.

**Figure 3:**
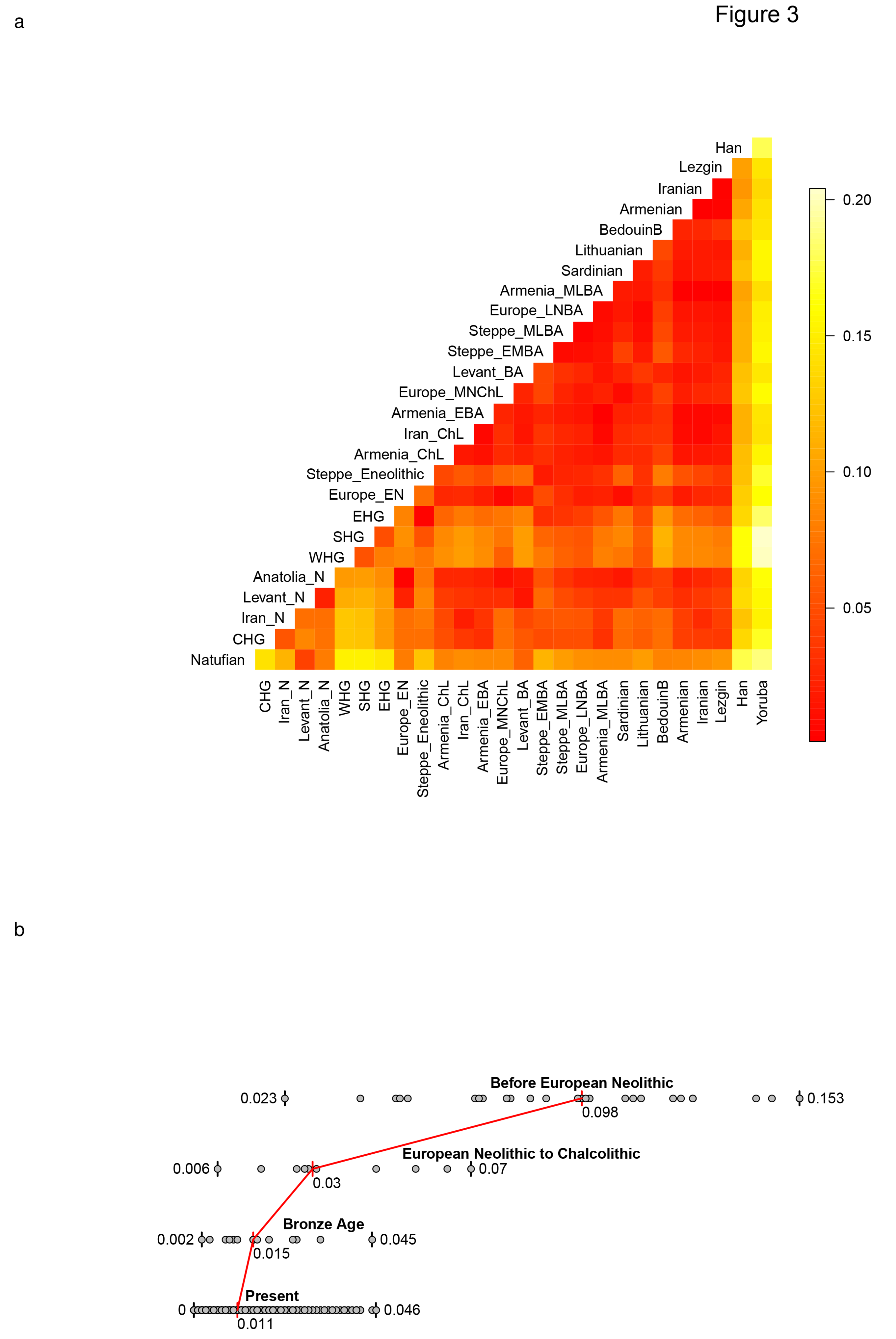
Genetic differentiation and its dramatic decrease over time in West Eurasia. (a) Pairwise F_ST_ between 19 Ancient West Eurasian populations (arranged in approximate chronological order), and select present-day populations. (b) Pairwise F_ST_ distribution among populations belonging to four successive time slices in West Eurasia; the median (red) and range of F_ST_ is shown.

## Continuity between pre-farming hunter-gatherers and early farmers of the Near East

Our data document continuity across the hunter-gatherer / farming transition, separately in the southern Levant and in the southern Caucasus-Iran highlands. The qualitative evidence for this is that PCA, ADMIXTURE, and outgroup *f*_3_ analysis cluster Levantine hunter-gatherers (Natufians) with Levantine farmers, and Iranian and Caucasus Hunter Gatherers with Iranian farmers (Fig. 1b; Extended Data Fig. 1; Extended Data Fig. 2). We confirm this in the Levant by showing that its early farmers share significantly more alleles with Natufians than with the early farmers of Iran: the statistic *f*_4_(Levant_N, Chimp; Natufian, Iran_N) is significantly positive (Z=13.6). The early farmers of the Caucasus-Iran highlands similarly share significantly more alleles with the hunter-gatherers of this region than with the early farmers from the Levant: the statistic *f*_4_(Iran_N, Chimp; Caucasus or Iran highland hunter-gatherers, Levant_N) is significantly positive (Z>6).

## How diverse first farmers of the Near East mixed to form the region’s later populations

Almost all ancient and present-day West Eurasians have evidence of significant admixture between two or more ancestral populations, as documented by statistics of the form *f*_3_(*Test; Reference*_1_, *Reference*_2_) which if negative, show that a *Test* population’s allele frequencies tend to be intermediate between two *Reference* populations^16^ (Extended Data Table 2). To better understand the admixture history beyond these patterns, we used *qpAdm*^1^, which can evaluate whether a particular *Test* population is consistent with being derived from a set of proposed source populations, and if so, infer mixture proportions (Methods). We used this approach to carry out a systematic survey of ancient West Eurasian populations to explore their possible sources of admixture (Fig. 4; Supplementary Information, section 7).

**Figure 4:**
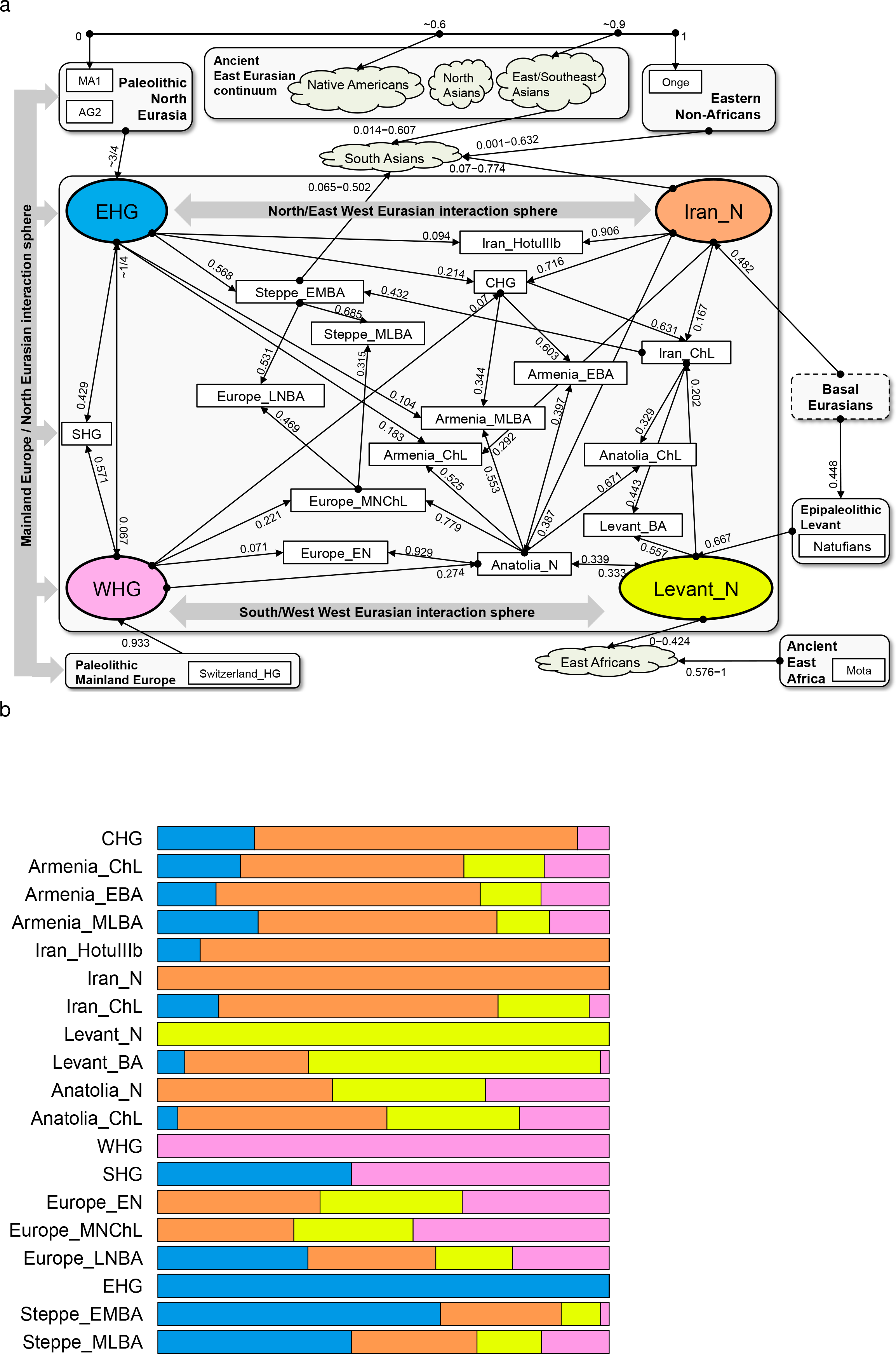
Modelling ancient West Eurasians, East Africans, East Eurasians and South Asians. (a) All the ancient populations can be modelled as mixtures of two or three other populations and up to four proximate sources (marked in colour). Mixture proportions inferred by *qpAdm* are indicated by the incoming arrows to each population. Clouds represent sets of more than one population. Multiple admixture solutions are consistent with the data for some populations, and while only one solution is shown here, Supplementary Information, section 7 presents the others. (b) A flat representation of the graph showing mixture proportions from the four proximate sources.

Among first farmers, those of the Levant trace ~2/3 of their ancestry to people related to Natufian hunter-gatherers and ~1/3 to people related to Anatolian farmers (Supplementary Information, section 7). Western Iranian first farmers cluster with the likely Mesolithic HotuIIIb individual and more remotely with hunter-gatherers from the southern Caucasus (Fig. 1b), and share alleles at an equal rate with Anatolian and Levantine early farmers (Supplementary Information, section 7), highlighting the long-term isolation of western Iran.

During subsequent millennia, the early farmer populations of the Near East expanded in all directions and mixed, as we can only model populations of the Chalcolithic and subsequent Bronze Age as having ancestry from two or more sources. The Chalcolithic people of western Iran can be modelled as a mixture of the Neolithic people of western Iran, the Levant, and Caucasus Hunter Gatherers (CHG), consistent with their position in the PCA (Fig. 1b). Admixture from populations related to the Chalcolithic people of western Iran had a wide impact, consistent with contributing ~44% of the ancestry of Levantine Bronze Age populations in the south and ~33% of the ancestry of the Chalcolithic northwest Anatolians in the west. Our analysis show that the ancient populations of the Chalcolithic Iran, Chalcolithic Armenia, Bronze Age Armenia and Chalcolithic Anatolia were all composed of the same ancestral components, albeit in slightly different proportions (Fig. 4b; Supplementary Information, section 7).

## The Near Eastern contribution to Europeans, East Africans and South Asians

Admixture did not only occur within the Near East but extended towards Europe. To the north, a population related to people of the Iran Chalcolithic contributed ~43% of the ancestry of early Bronze Age populations of the steppe. The spread of Near Eastern ancestry into the Eurasian steppe was previously inferred^7^ without access to ancient samples, by hypothesizing a population related to present-day Armenians as a source^7,8^. To the west, the early farmers of mainland Europe were descended from a population related to Neolithic northwestern Anatolians^8^. This is consistent with an Anatolian origin of farming in Europe, but does not reject other sources, since the spatial distribution of the Anatolian/European-like farmer populations is unknown. We can rule out the hypothesis that European farmers stem directly from a population related to the ancient farmers of the southern Levant^30,31^, however, since they share more allele with Anatolian Neolithic farmers than with Levantine farmers as attested by the positive statistic *f*_4_(Europe_EN, Chimp; Anatolia_N, Levant_N) (Z=15).

Migrations from the Near East also occurred towards the southwest into East African populations which experienced West Eurasian admixture ~1,000 BCE^32,33^. Previously, the West Eurasian population known to be the best proxy for this ancestry was present-day Sardinians ^33^, who resemble Neolithic Europeans genetically ^13,34^. However, our analysis shows that East African ancestry is significantly better modelled by Levantine early farmers than by Anatolian or early European farmers, implying that the spread of this ancestry to East Africa was not from the same group that spread Near Eastern ancestry into Europe (Extended Data Fig. 4; Supplementary Information, section 8).

In South Asia, our dataset provides insight into the sources of Ancestral North Indians (ANI), a West Eurasian related population that no longer exists in unmixed form but contributes a variable amount of the ancestry of South Asians^35,36^ (Supplementary Information, section 9) (Extended Data Fig. 4). We show that it is impossible to model the ANI as being derived from any single ancient population in our dataset. However, it can be modelled as a mix of ancestry related to both early farmers of western Iran and to people of the Bronze Age Eurasian steppe; all sampled South Asian groups are inferred to have significant amounts of both ancestral types. The demographic impact of steppe related populations on South Asia was substantial, as the Mala, a south Indian population with minimal ANI along the ‘Indian Cline’ of such ancestry^35,36^ is inferred to have ~18% steppe-related ancestry, while the Kalash of Pakistan are inferred to have ~50%, similar to present-day northern Europeans^7^.

## Broader insights into population transformations across West Eurasia and beyond

We were concerned that our conclusions might be biased by the particular populations we happened to sample, and that we would have obtained qualitatively different conclusions without data from some key populations. We tested our conclusions by plotting the inferred position of admixed populations in PCA against a weighted combination of their inferred source populations and obtained qualitatively consistent results (Extended Data Fig. 5).

To further assess the robustness of our inferences, we developed a method to infer the existence and genetic affinities of ancient populations from unobserved ‘ghost’ populations (Supplementary Information, section 10; Extended Data Fig. 6). This method takes advantage of the insight that if an unsampled ghost population admixes with differentiated ‘substratum’ populations, it is possible to extrapolate its identity by intersecting clines of populations with variable proportions of ‘ghost’ and ‘substratum’ ancestry. Applying this while withholding major populations, we validated some of our key inferences, successfully inferring mixture proportions consistent with those obtained when the populations are included in the analysis. Application of this methods highlights the impact of Ancient North Eurasian (ANE) ancestry related to the ~22,000 BCE Mal’ta 1 and ~15,000 BCE Afontova Gora 2^15^ on populations living in Europe, the Americas, and Eastern Eurasia. Eastern Eurasians can be modelled as arrayed along a cline with different proportions of ANE ancestry (Supplementary Information, section 11; Extended Data Fig. 7), ranging from ~40% ANE in Native Americans matching previous findings^13,15^, to no less than ~5-10% ANE in diverse East Asian groups including Han Chinese (Extended Data Fig. 4; Extended Data Fig. 6f). We also document a cline of ANE ancestry across the east-west extent of Eurasia. Eastern Hunter Gatherers (EHG) derive ~3/4 of their ancestry from the ANE (Supplementary Information, section 11); Scandinavian hunter-gatherers^7,8,13^ (SHG) are a mix of EHG and WHG; and WHG are a mix of EHG and the Upper Paleolithic Bichon from Switzerland (Supplementary Information, section 7). Northwest Anatolians—with ancestry from a population related to European hunter-gatherers (Supplementary Information, section 7)—are better modelled if this ancestry is taken as more extreme than Bichon (Supplementary Information, section 10).

The population structure of the ancient Near East was not independent of that of Europe (Supplementary Information, section 4), as evidenced by the highly significant (Z=‑8.9) statistic *f*_4_(Iran_N, Natufian;WHG, EHG) which suggests gene flow in ‘northeastern’ (Neolithic Iran/EHG) and ‘southwestern’ (Levant/WHG) interaction spheres (Fig. 4d). This interdependence of the ancestry of Europe and the Near East may have been mediated by unsampled geographically intermediate populations^37^ that contribute ancestry to both regions.

## Conclusions

By analysing genome-wide ancient DNA data from ancient individuals from the Levant, Anatolia, the southern Caucasus and Iran, we have provided a first glimpse of the demographic structure of the human populations that transitioned to farming. We reject the hypothesis that the spread of agriculture in the Near East was achieved by the dispersal of a single farming population displacing the hunter-gatherers they encountered. Instead, the spread of ideas and farming technology moved faster than the spread of people, as we can determine from the fact that the population structure of the Near East was maintained throughout the transition to agriculture. A priority for future ancient DNA studies should be to obtain data from older periods, which would reveal the deeper origins of the population structure in the Near East. It will also be important to obtain data from the ancient civilizations of the Near East to bridge the gap between the region’s prehistoric inhabitants and those of the present.

## Acknowledgements

We thank the 238 human subjects who voluntarily donated the samples whose genome-wide data we newly report in this study. We thank D. Labuda for sharing the collection of DNA samples from Poland, and P. Zalloua for sharing the collection of DNA samples from Lebanon. We thank O. Bar-Yosef, M. Bonogofsky, I. Hershkowitz, M. Lipson, I. Mathieson, H. May, R. Meadow, I. Olalde, S. Paabo, P. Skoglund, and N. Nakatsuka for comments and critiques, and M. Ferry and M. Michel for their work on the in-solution enrichment experiments. S.C. was supported by the Irish Research Council for Humanities and Social Sciences (IRCHSS) ERC Support Programme. Q.F. was funded by the Bureau of International Cooperation of Chinese Academy of Sciences, the National Natural Science Foundation of China (L1524016) and the Chinese Academy of Sciences Discipline Development Strategy Project (2015-DX-C-03). The Scottish diversity data from Generation Scotland received funding from the Chief Scientist Office of the Scottish Government Health Directorates [CZD/16/6] and the Scottish Funding Council [HR03006], while the Scottish Donor DNA Databank (GS:3D) was funded by a project grant from the Scottish Executive Health Department, Chief Scientist Office [CZB/4/285]. M.S., A.Tön., M.B. and P.K. were supported by grants from the Collaborative Research Center funded by the German Research Foundation (CRC 1052; B01, B03, C01). M.S.-P. was partially funded by a Wenner-Gren Foundation Dissertation Fieldwork Grant (#9005), and by the National Science Foundation DDRIG (BCS-1455744). P.K. was supported by the Federal Ministry of Education and Research (BMBF), Germany (FKZ: 01EO1501). J.F.W acknowledges the MRC “QTL in Health and Disease” programme grant. The Romanian contribution to this work was supported by the EC Commission, Directorate General XII, within the framework of the Cooperation in Science and Technology with Central and Eastern European Countries (Supplementary Agreement ERBCIPDCT 940038 to the Contract ERBCHRXCT 920032, coordinated by Prof. A. Piazza, Turin, Italy). M.R. received support from the Leverhulme Trust’s Doctoral Scholarship programme. O.S. and A.Tor. were supported by the University of Pavia strategic theme “Towards a governance model for international migration: an interdisciplinary and diachronic perspective” (MIGRAT-IN-G) and the Italian Ministry of Education, University and Research: Progetti Ricerca Interesse Nazionale 2012. The Raqefet Cave Natufian project was supported by funds from the National Geographic Society (Grant #8915-11), the Wenner-Gren Foundation (Grant #7481) and the Irene Levi-Sala CARE Foundation, while radiocarbon dating on the samples was funded by the Israel Science Foundation (Grant 475/10; E. Boaretto). R.P. was supported by ERC starting grant ADNABIOARC (263441). D R. was supported by NIH grant GM100233, by NSF HOMINID BCS-1032255, and is a Howard Hughes Medical Institute investigator.

## Author Contributions

R.P. and D.R. conceived the idea for the study. D.N., G.R., D.C.M., S.C., S.A., G.L., F.B., B.Gas., J.M.M., M.G., V.E., A.M., C.M., F.G., N.A.H. and R.P. assembled archaeological material. N.R., D.F., M.N., B.Gam., K.Si., S.C., K.St., E.H., Q.F., G.G.-F., R.P. and D.R. performed or supervised ancient DNA wet laboratory work. L.B, M.B., A.C., G.C., D.C., P.F., E.G., S.M.K., P.K., J.K., D M., M.M., D.A.M., S O., M R., O S., M.S.-P., G.S., M.S., A.Ton., A.Tor., J.F.W., L.Y. and D.R. assembled present-day samples for genotyping. I.L, N.P. and D.R. developed methods for data analysis. I.L., S.M., Q.F., N.P. and D.R. analyzed data. I.L., R.P. and D.R. wrote the manuscript and supplements. All authors read the manuscript and provided comments.

## Author Information

The aligned sequences are available through the European Nucleotide Archive under accession number xxx. Fully public subsets of the analysis datasets are at (http://genetics.med.harvard.edu/reichlab/Reich_Lab/Datasets.html). The complete dataset (including present-day humans for which the informed consent is not consistent with public posting of data) is available to researchers who send a signed letter to D.R. indicating that they will abide by specified usage conditions (Supplementary Information, section 2).

## Online Methods

### Ancient DNA data

In a dedicated ancient DNA laboratory at University College Dublin, we prepared powder from 132 ancient Near Eastern samples, either by dissecting the inner ear region of the petrous bone using a sandblaster (Renfert), or by drilling using a Dremel tool and single-use drill bits and selecting the best preserved bone fragments based on anatomical criteria. These fragments were then powdered using a mixer mill (Retsch Mixer Mill 400)^4^.

We performed all subsequent processing steps in a dedicated ancient DNA laboratory at Harvard Medical School, where we extracted DNA from the powder (usually 75 mg, range 14-81 mg) using an optimized ancient DNA extraction protocol^38^, but replaced the assembly of Qiagen MinElute columns and extension reservoirs from Zymo Research with a High Pure Extender Assembly from the High Pure Viral Nucleic Acid Large Volume Kit (Roche Applied Science). We built a total of 170 barcoded double-stranded Illumina sequencing libraries for these samples^39^, of which we treated 167 with Uracil-DNA glycosylase (UDG) to remove the characteristic C-to-T errors of ancient DNA^40^. The UDG treatment strategy is (by-design) inefficient at removing terminal uracils, allowing the mismatch rate to the human genome at the terminal nucleotide to be used for authentication^39^. We updated this library preparation protocol in two ways compared to the original publication: first, we used 16U Bst2.0 Polymerase, Large Fragment (NEB) and 1x Isothermal Amplification buffer (NEB) in a final volume of 25 μL fill-in reaction, and second, we used the entire inactivated 25μL fill-in reaction in a total volume of 100μL PCR mix with 1 μM of each primer^41^. We included extraction negative controls (where no sample powder was used) and library negative controls (where extract was supplemented by water) in every batch of samples processed and carried them through the entire wet lab processing to test for reagent contamination.

We screened the libraries by hybridizing them in solution to a set of oligonucleotide probes tiling the mitochondrial genome^42^, using the protocol described previously^7^. We sequenced the enriched libraries using an Illumina NextSeq 500 instrument using 2×76bp reads, trimmed identifying sequences (seven base pair molecular barcodes at either end) and any trailing adapters, merged read pairs that overlapped by at least 15 base pairs, and mapped the merged sequences to the RSRS mitochondrial DNA reference genome^43^, using the Burrows Wheeler Aligner^44^ *(bwa)* and the command *samse* (v0.6.1).

We enriched promising libraries for a targeted set of ~1.2 million SNPs^8^ as in ref. 5, and adjusted the blocking oligonucleotide and primers to be appropriate for our libraries. The specific probe sequences are given in Supplementary Data 2 of ref. 7 (http://www.nature.com/nature/journal/v522/n7555/abs/nature14317.html#supplementary-information) and Supplementary Data 1 of ref. 6. (http://www.nature.com/nature/journal/v524/n7564/full/nature14558.html#supplementary-information). We sequenced the libraries on an Illumina NextSeq 500 using 2×76bp reads. We trimmed identifying sequences (molecular barcodes) and any trailing adapters, merged pairs that overlapped by at least 15 base pairs (allowing up to one mismatch), and mapped the merged sequences to *hg19* using the single-ended aligner *samse* in bwa (v0.6.1). We removed duplicated sequences by identifying sets of sequences with the same orientation and start and end positions after alignment to *hg19;* we picked the highest quality sequence to represent each set. For each sample, we represented each SNP position by a randomly chosen sequence, restricting to sequences with a minimum mapping quality (MAPQ≥10), sites with a minimum sequencing quality (≥20), and removing 2 bases at the ends of reads. We sequenced the enriched products up to the point that we estimated that generating a hundred new sequences was expected to add data on less than about one new SNP^8^.

### Testing for contamination and quality control

For each ancient DNA library, we evaluated authenticity in several ways. First, we estimated the rate of matching to the consensus sequence for mitochondrial genomes sequenced to a coverage of at least 10-fold from the initial screening data. Of the 76 libraries that contributed to our dataset (coming from 45 samples), 70 had an estimated rate of sequencing matching to the consensus of >95% according to contamMix^5^ (the remaining libraries had estimated match rates of 75-92%, but gave no sign of being outliers in principal component analysis or X chromosome contamination analysis so we retained them for analysis) (Supplementary Data Table 1). We quantified the rate of C-to-T substitution in the final nucleotide of the sequences analyzed, relative to the human reference genome sequence, and found that all the libraries analyzed had rates of at least 3%^39^, consistent with genuine ancient DNA. For the nuclear data from males, we used the ANGSD software^45^ to estimate a conservative X chromosome estimate of contamination. We determined that all libraries passing our quality control and for which we had sufficient X chromosome data to make an assessment had contamination rates of 0-1.5%. Finally, we merged data for samples for which we had multiple libraries to produce an analysis dataset.

### Affymetrix Human Origins genotyping data

We genotyped 238 present-day individuals from 17 diverse West Eurasian populations on the Affymetrix Human Origins array^16^, and applied quality control analyses as previously described^13^ (Supplementary Data Table 2). We merged the newly generated data with data from 2,345 individuals previously genotyped on the same array^13^. All individuals that were genotyped provided informed consent consistent with studies of population history, following protocols approved by the ethical review committees of the institutions of the researchers who collected the samples. De-identified aliquots of DNA from all individuals were sent to the core facility of the Center for Applied Genomics at the Children’s Hospital of Philadelphia for genotyping and data processing. For 127 of the individuals with newly reported data, the informed consent was consistent with public distribution of data, and the data can be downloaded at http://genetics.med.harvard.edu/reich/Reich_Lab/Datasets.html. To access data for the remaining 111 samples, researchers should a signed letter to D.R. containing the following text: “(a) I will not distribute the samples marked “signed letter” outside my collaboration; (b) I will not post data from the samples marked “signed letter” publicly; (c) I will make no attempt to connect the genetic data for the samples marked “signed letter” to personal identifiers; (d) I will not use the data for samples marked “signed letter” for commercial purposes.” Supplementary Data Table 2 specifies which samples are consistent with which type of data distribution.

### Datasets

We carried out population genetic analysis on two datasets: (i) *HO* includes 2,583 present-day humans genotyped on the Human Origins array^13,16^ including 238 newly reported (Supplementary Data Table 2; Supplementary Information, section 2), and 281 ancient individuals on a total of 592,146 autosomal SNPs. (ii) *HOIll* includes the 281 ancient individuals on a total of 1,055,186 autosomal SNPs, including those present in both the Human Origins and Illumina genotyping platforms, but excluding SNPs on the sex chromosomes or additional SNPs of the 1240k capture array that were included because of their potential functional importance^8^. We used *HO* for analyses that involve both ancient and present-day individuals, and *HOIll* for analysis on ancient individuals alone. We also use 235 individuals from Pagani et al.^32^ on 418,700 autosomal SNPs to study admixture in East Africans (Supplementary Information, section 8). Ancient individuals are represented in ‘pseudo-haploid’ form by randomly choosing one allele for each position of the array.

### Principal Components Analysis

We carried out principal components analysis in the *smartpca* program of EIGENSOFT^17^, using default parameters and the lsqproject: YES^13^ and numoutlieriter: 0 options. We carried out PCA of the *HO* dataset on 991 present-day West Eurasians (Extended Data Fig. 1), and projected the 278 ancient individuals (Fig. 1b).

### ADMIXTURE Analysis

We carried out ADMIXTURE analysis^18^ of the *HO* dataset after pruning for linkage disequilibrium in PLINK^46,47^ with parameters--indep-pairwise 200 25 0.4 which retained 296,309 SNPs. We performed analysis in 20 replicates with different random seeds, and retained the highest likelihood replicate for each value of K. We show the K=11 results for the 281 ancient samples in Fig. 1c (this is the lowest K for which components maximized in European hunter-gatherers, ancient Levant, and ancient Iran appear).

### *f*-statistics

We carried out analysis of *f*_3_-statistics, *f*_4_-ratio, and *f*_4_-statistics statistics using the ADMIXTOOLS^16^ programs *qp3Pop, qpF4ratio* with default parameters, and *qpDstat* with f4mode: YES, and computed standard errors with a block jackknife^48^. For computing *f*_3_-statistics with an ancient population as a target, we set the inbreed:YES parameter. We computed *f*-statistics on the *HOIll* dataset when no present-day humans were involved and on the *HO* dataset when they were. We computed the statistic *f*_4_(*Test*, Mbuti; Altai, Denisovan) in Fig. 2 on the *HOIll* dataset after merging with whole genome data on 3 Mbuti individuals from Panel C of the Simons Genome Diversity Project^49^. We computed the dendrogram of Extended Data Fig. 2 showing hierarchical clustering of populations with outgroup *f*_3_-statistics using the open source *heatmap.2* function of the *gplots* package in *R*.

### Negative correlation of Basal Eurasian ancestry with Neanderthal ancestry

We used the *lm* function of R to fit a linear regression of the rate of allele sharing of a *Test* population with the Altai Neanderthal as measured by *f*_4_(Test, Mbuti; Altai, Denisovan) as the dependent variable, and the proportion of Basal Eurasian ancestry (Supplementary Information, section 4) as the predictor variable,. Extrapolating from the fitted line, we obtain the value of the statistic expected if *Test* is a population of 0% or 100% Basal Eurasian ancestry. We then compute the ratio of the Neanderthal ancestry estimate in Basal Eurasians relative to non-Basal Eurasians as74(100% Basal Eurasian, Mbuti; Altai, Denisovan)/f4(0% Basal Eurasian, Mbuti; Altai, Denisovan). We use a block jackknife^48^, dropping one of 100 contiguous blocks of the genome at a time, to estimate the value and standard error of this quantity (9±26%). We compute a 95% confidence interval based on the point estimate ±1.96-times the standard error: −42 to 60%. We truncated to 0-60% on the assumption that Basal Eurasians had no less Neanderthal admixture than Mbuti from sub-Saharan Africa.

### Estimation of F_ST_ coefficients

We estimated F_ST_ in *smartpca*^17^ with default parameters, inbreed: YES, and fstonly: YES.

### Admixture Graph modeling

We carried out Admixture Graph modeling with the *qpGraph* software^16^ using Mbuti as an outgroup unless otherwise specified.

### Testing for the number of streams of ancestry

We used the *qpWave*^35,50^ software, described in Supplementary Information, section 10 of ref.^7^, to test whether a set of ‘Left’ populations is consistent with being related via as few as *N* streams of ancestry to a set of ‘Right’ populations by studying statistics of the form *X*(*u*, *v*) = *F*_4_(*u*_0_, *u*; *v*_0_, *v*) where *u*_0_, *v*_0_ are basis populations chosen from the ‘Left’ and ‘Right’ sets and u, v are other populations from these sets. We use a Hotelling’s T^2^ test^50^ to evaluate whether the matrix of size (L−1)*(R−1), where L, R are the sizes of the ‘Left’ and ‘Right’ sets has rank *m*. If this is the case, we can conclude that the ‘Left’ set is related via at least *N*=*m*+1 streams of ancestry related differently to the ‘Right’ set.

### Inferring mixture proportions without an explicit phylogeny

We used the *qpAdm* methodology described in Supplementary Information, section 10 of ref. ^7^ to estimate the proportions of ancestry in a *Test* population deriving from a mixture of *N* ‘reference’ populations by exploiting (but not explicitly modeling) shared genetic drift with a set of ‘Outgroup’ populations (Supplementary Information, section 7). We set the details: YES parameter, which reports a normally distributed Z-score estimated with a block jackknife for the difference between the statistics *f*_4_(*u*_0_, *Test*; *v*_0_, *v*) and*f*_4_(*u*_0_, *Estimated Test; v*_0_, *v*) where *Estimated Test* is 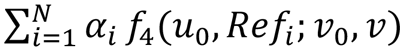 the average of these *f*_4_-statistics weighed by the mixture proportions *α*_*i*_ from the *N* reference populations.

### Modeling admixture from ghost populations

We model admixture from a ‘ghost’ (unobserved) population *X* in the specific case that *X* has part of its ancestry from two unobserved ancestral populations *p* and *q.* Any population *X* composed of the same populations *p* and *q* resides on a line defined by two observed reference populations *r*_1_ and *r*_2_ composed of the same elements *p* and *q* according to a parametric equation *x* = *r*_1_ + *λ*(*r*_2_ — *r*_1_) with real-valued parameter *λ*. We define and solve the optimization problem of fitting *λ* and obtain mixture proportions (Supplementary Information, section 10).

## Extended Data Tables and Extended Data Figure Legends

**Extended Data Table 1:** No evidence for admixture related to sub-Saharan Africans in Natufians. We computed the statistic *f*_4_(*Natufian*, *Other Ancient*; *African*, Chimp) varying *African* to be Mbuti, Yoruba, Ju_hoan_North, or the ancient Mota individual. Gene flow between Natufians and African populations would be expected to bias these statistics positive. However, we find most of them to be negative in sign and all of them to be non-significant (|Z|<3), providing no evidence that Natufians differ from other ancient samples with respect to African populations.

| *Other Ancient* | *African* | $f_4(\text{Natufian}, \text{Other Ancient}, \text{African}, \text{Chimp})$ | *Z* | Number of SNPs |
| --- | --- | --- | --- | --- |
| EHG | Mbuti | -0.00044 | -1.0 | 254033 |
| EHG | Yoruba | 0.00029 | 0.7 | 254033 |
| EHG | Ju_hoan_North | -0.00015 | -0.4 | 254033 |
| EHG | Mota | -0.00022 | -0.4 | 253986 |
| WHG | Mbuti | -0.00067 | -1.7 | 261514 |
| WHG | Yoruba | -0.00045 | -1.1 | 261514 |
| WHG | Ju_hoan_North | -0.00046 | -1.2 | 261514 |
| WHG | Mota | -0.00129 | -2.3 | 261461 |
| SHG | Mbuti | -0.00076 | -2.0 | 255686 |
| SHG | Yoruba | -0.00039 | -1.0 | 255686 |
| SHG | Ju_hoan_North | -0.00052 | -1.4 | 255686 |
| SHG | Mota | -0.00091 | -1.7 | 255641 |
| Switzerland_HG | Mbuti | -0.00018 | -0.4 | 261322 |
| Switzerland_HG | Yoruba | 0.00019 | 0.4 | 261322 |
| Switzerland_HG | Ju_hoan_North | 0.00009 | 0.2 | 261322 |
| Switzerland_HG | Mota | -0.00062 | -0.9 | 261276 |
| Kostenki14 | Mbuti | 0.00034 | 0.7 | 246765 |
| Kostenki14 | Yoruba | 0.00120 | 2.3 | 246765 |
| Kostenki14 | Ju_hoan_North | 0.00069 | 1.4 | 246765 |
| Kostenki14 | Mota | 0.00036 | 0.5 | 246719 |
| MA1 | Mbuti | -0.00038 | -0.7 | 191819 |
| MA1 | Yoruba | 0.00009 | 0.2 | 191819 |
| MA1 | Ju_hoan_North | -0.00010 | -0.2 | 191819 |
| MA1 | Mota | -0.00038 | -0.5 | 191782 |
| CHG | Mbuti | -0.00051 | -1.2 | 261505 |
| CHG | Yoruba | -0.00012 | -0.3 | 261505 |
| CHG | Ju_hoan_North | -0.00013 | -0.3 | 261505 |
| CHG | Mota | -0.00042 | -0.7 | 261456 |
| Iran_N | Mbuti | -0.00018 | -0.4 | 232927 |
| Iran_N | Yoruba | 0.00036 | 0.8 | 232927 |
| Iran_N | Ju_hoan_North | 0.00041 | 0.9 | 232927 |
| Iran_N | Mota | 0.00006 | 0.1 | 232880 |

**Extended Data Table 2:**
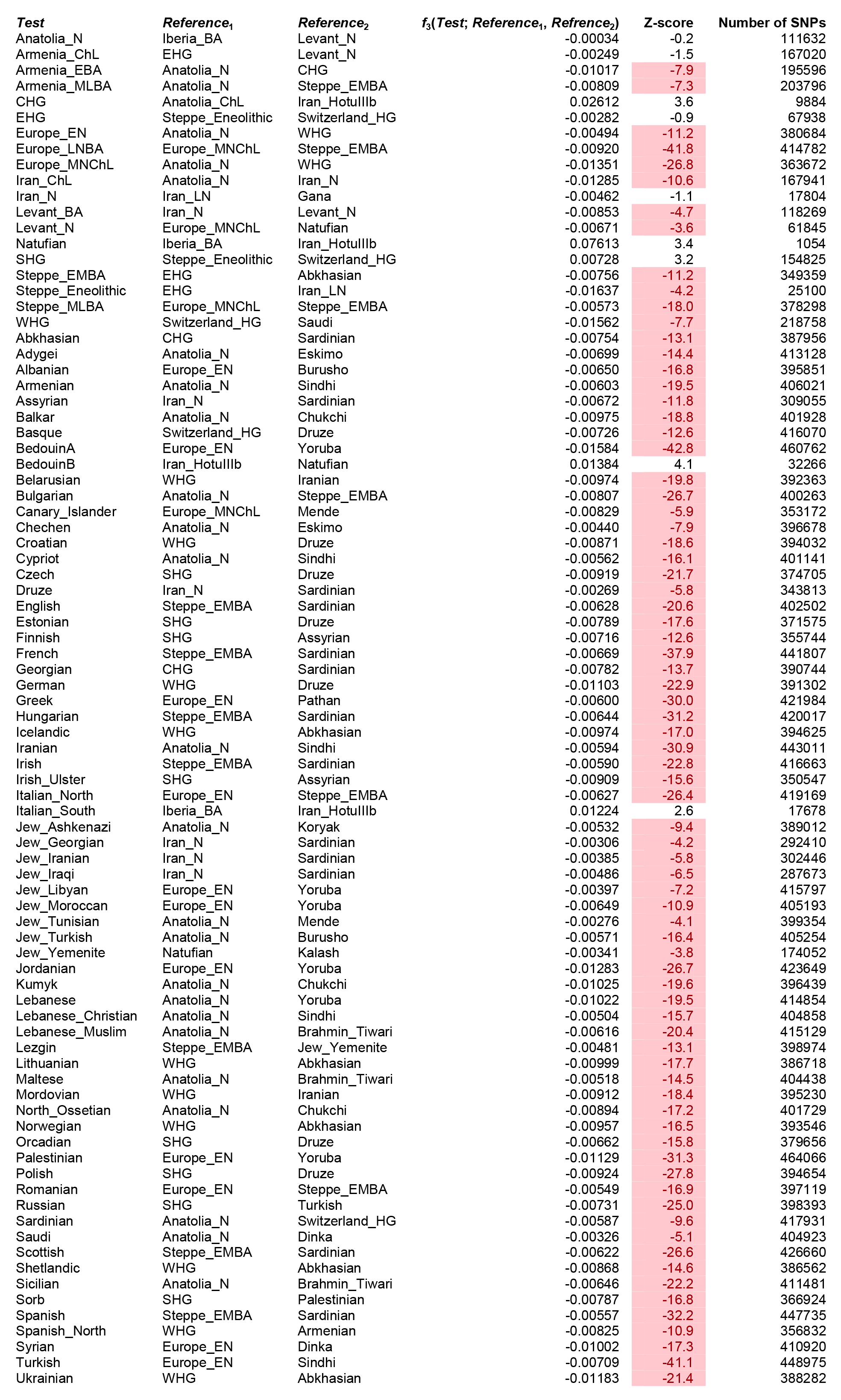
Admixture *f*_3_-statistics. We show the lowest Z-score of the statistic *f*_3_(*Test*; *Reference*_1_, *Refrence*_2_) for every ancient *Test* population with at least 2 individuals and every pair (*Reference*_1_, *Refrence*_2_) of ancient or present-day source populations. Z-scores lower than-3 are highlighted and indicate that the *Test* population is admixed from sources related to (but not identical to) the reference populations. Z-scores greater than-3 are consistent with the population either being admixed or not.

**Extended Data Figure 1:**
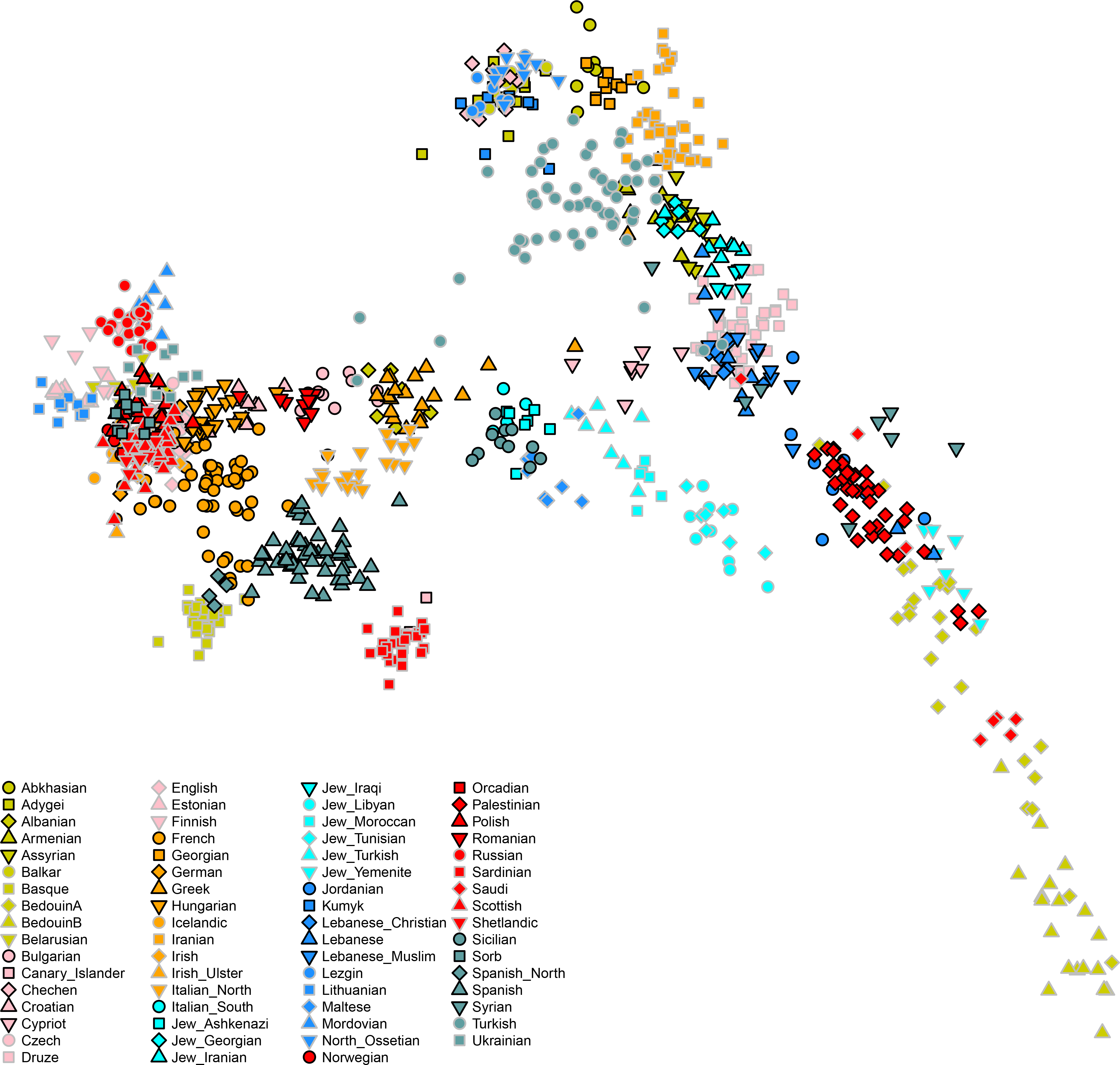
Principal components analysis of 991 present-day West Eurasians. The PCA analysis is performed on the same set of individuals as are reported in Fig. 1b, using EIGENSOFT. Here, we color the samples by population (to highlight the present-day populations) instead of using grey points as in Fig. 1b (where the goal is to highlight ancient samples).

**Extended Data Figure 2:**
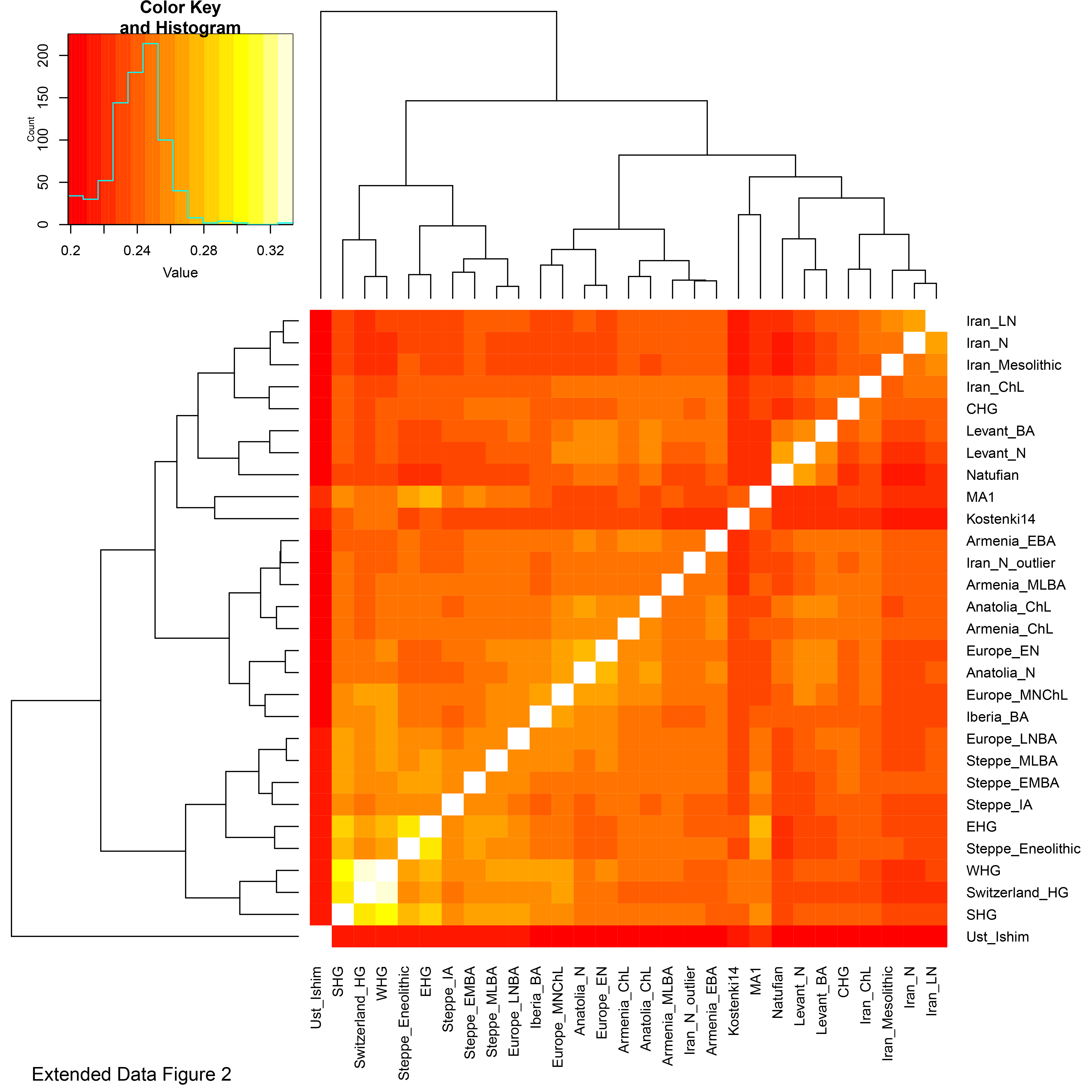
Outgroup *f*_3_(Mbuti; X, Y) for pairs of ancient populations. The dendrogram is plotted for convenience and should not be interpreted as a phylogenetic tree. Areas of high shared genetic drift are ‘yellow’ and include from top-right to bottom-left along the diagonal: early Anatolian and European farmers; European hunter-gatherers, Steppe populations and ones admixed with steppe ancestry; populations from the Levant from the Epipaleolithic (Natufians) to the Bronze Age; populations from Iran from the Mesolithic to the Late Neolithic.

**Extended Data Figure 3:**
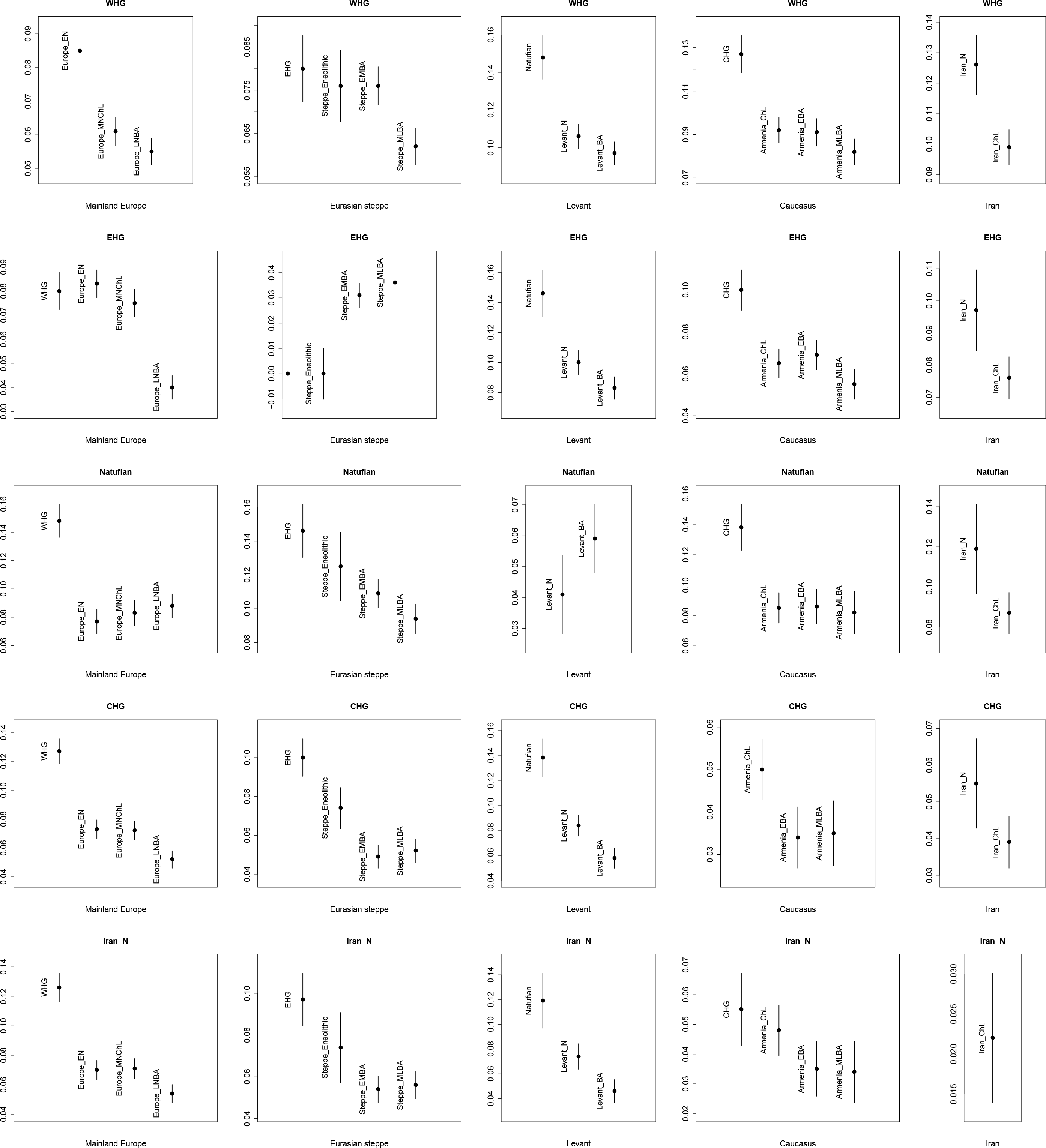
Reduction of genetic differentiation in West Eurasia over time. We measure differentiation by F_ST_. Each column of the 5x5 matrix of plots represents a major region and each row the earliest population with at least two individuals from each major region.

**Extended Data Figure 4:**
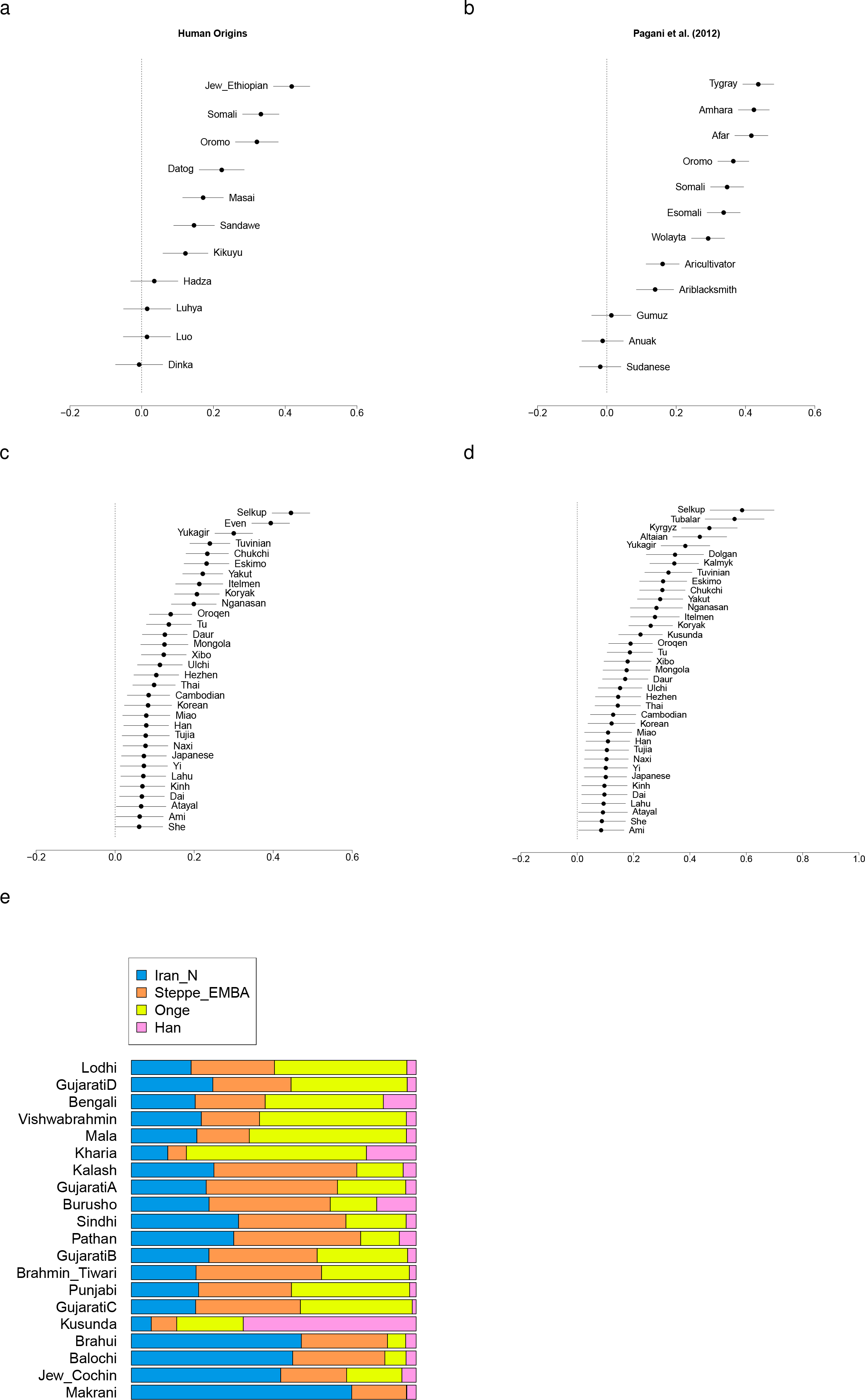
West Eurasian related admixture in East Africa, Eastern Eurasia and South Asia. (a) Levantine ancestry in Eastern Africa in the Human Origins dataset, (b) Levantine ancestry in different Eastern African population in the dataset of Pagani et al. (2012); the remainder of the ancestry is a clade with Mota, a ~4,500 year old sample from Ethiopia. (c) EHG ancestry in Eastern Eurasians, or (d) Afontova Gora (AG2) ancestry in Eastern Eurasians; the remainder of their ancestry is a clade with Onge. (e) Mixture proportions for South Asian populations showing that they can be modelled as having West Eurasian-related ancestry similar to that in populations from both the Eurasian steppe and Iran.

**Extended Data Figure 5:**
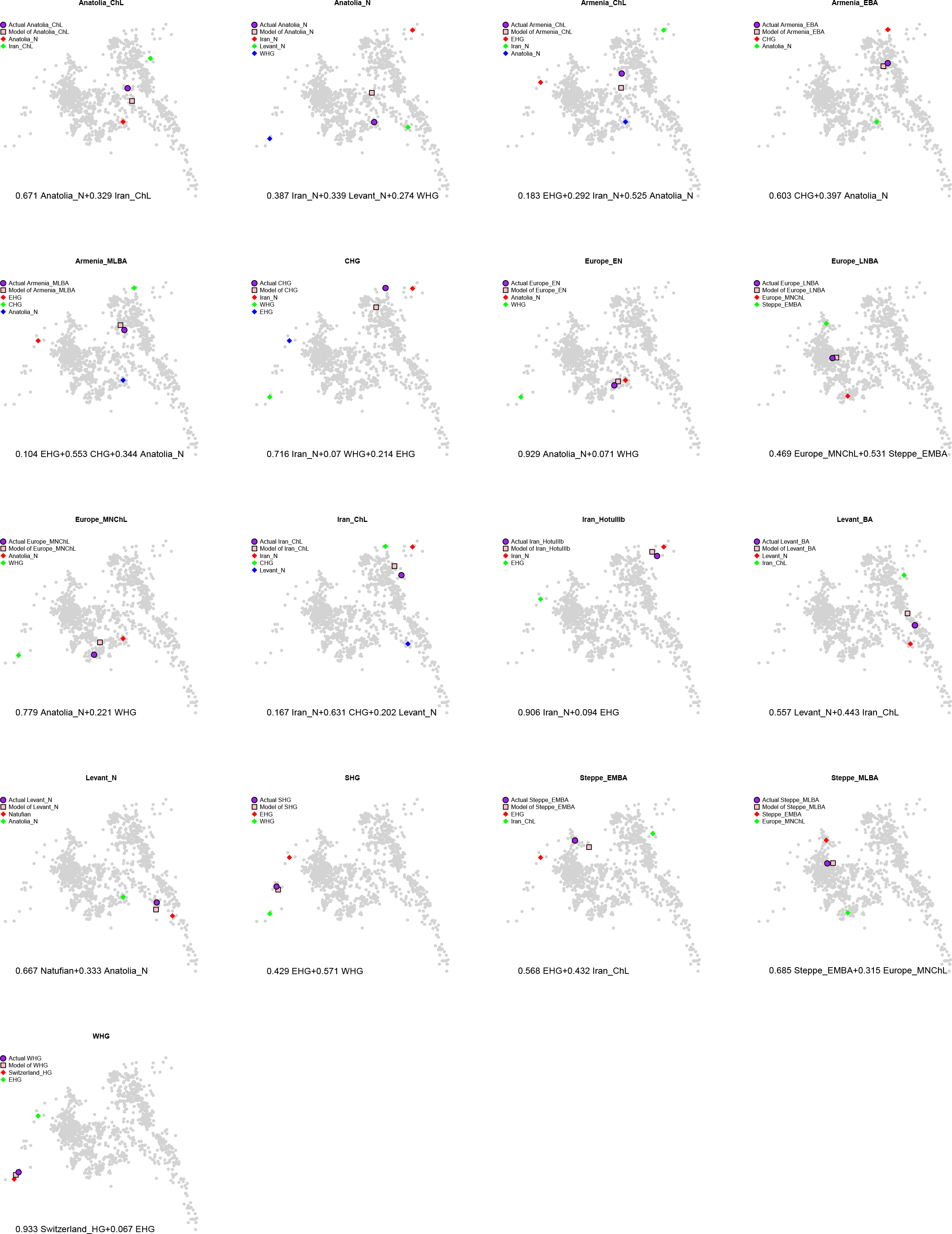
**Inferred position of ancient populations in West Eurasian PCA according to the model of Fig. 4**.

**Extended Data Figure 6:**
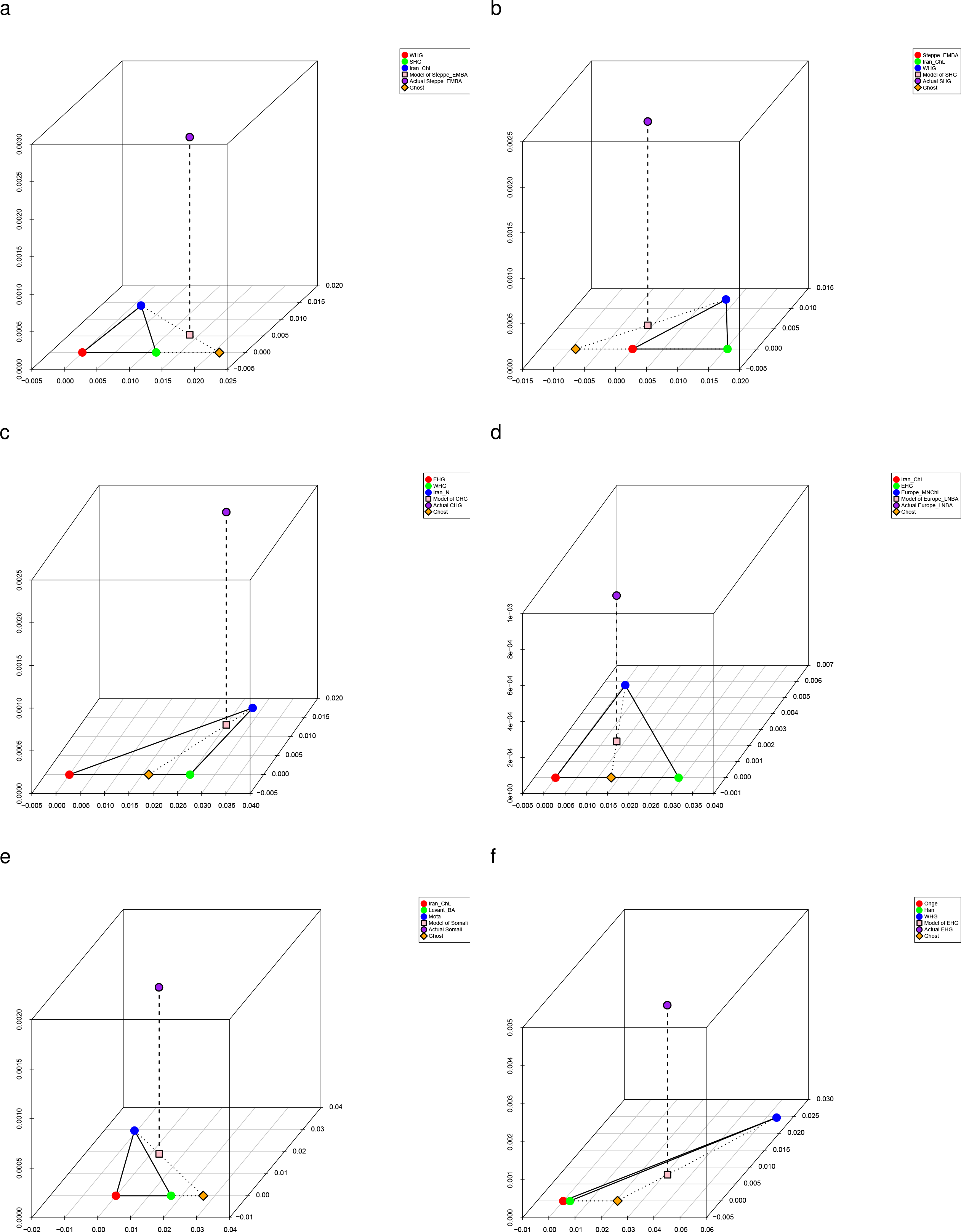
Admixture from ghost populations using ‘cline intersection’. We model each *Test* population (purple) in panels (a-f) as a mixture (pink) of a fixed reference population (blue) and a ghost population (orange) residing on the cline defined by two other populations (red and green) according to the visualization method of Supplementary Information, section 10. (a) Early/Middle Bronze Age steppe populations are a mixture of Iran_ChL and a population on the WHG→SHG cline. (b) Scandinavian hunter-gatherers (SHG) are a mixture of WHG and a population on the Iran_ChL→Steppe_EMBA cline. (c) Caucasus hunter-gatherers (CHG) are a mixture of Iran_N and both WHG and EHG. (d) Late Neolithic/Bronze Age Europeans are a mixture of the preceding Europe_MNChL population and a population with both EHG and Iran_ChL ancestry. (e) Somali are a mixture of Mota and a population on the Iran_ChL→Levant_BA cline. (f) Eastern European hunter-gatherers (EHG) are a mixture of WHG and a population on the Onge → Han cline.

**Extended Data Figure 7:**
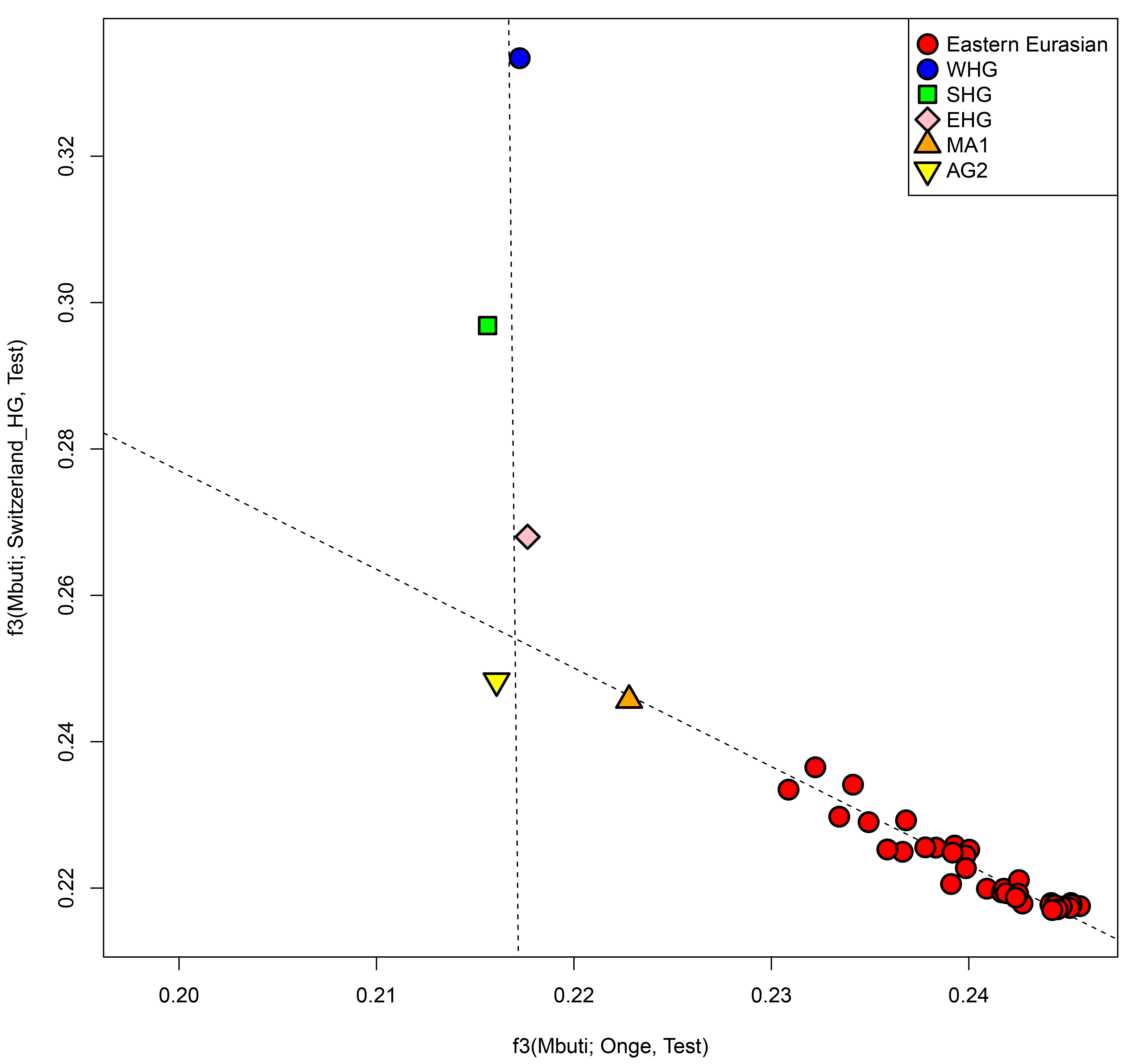
Admixture from a ‘ghost’ ANE population into both European and Eastern Eurasian ancestry. EHG, and Upper Paleolithic Siberians Mal’ta 1 (MA1) and Afontova Gora 2 (AG2) are positioned near the intersection of clines formed by European hunter-gatherers (WHG, SHG, EHG) and Eastern non-Africans in the space of outgroup *f*_3_-statistics of the form *f*_3_(Mbuti; Papuan, *Test)* and *f*_3_(Mbuti; Switzerland_HG, *Test*).

